# Cell cycle-regulated tug-of-war between microtubule motors positions major trafficking organelles

**DOI:** 10.1101/2025.10.01.679872

**Authors:** Avishkar V. Sawant, Irina Kaverina

**Affiliations:** Department of Cell and Developmental Biology, Vanderbilt University, Nashville, TN, USA

**Keywords:** Golgi, ER exit sites, centrosomes, CDK1, Kinesin-1, KIF5B, KLC, Dynein, KIFC3, cell cycle

## Abstract

The morphology and integrity of the Golgi apparatus, which are critical for protein processing and sorting, undergo significant changes during interphase in proliferating mammalian cells. Initially compact during G1, the Golgi dissociates from the centrosomes in S-phase and transitions into an extended perinuclear ribbon in G2. The mechanisms underlying these changes remain unclear. Golgi positioning can be influenced by microtubule (MT)-dependent motors and membrane exchange with endoplasmic reticulum exit sites (ERES). We employ loss-of-function approaches and live-cell microscopy to demonstrate that CDK1-dependent signaling modulates the tug-of-war between plus- and minus-end-directed motors, resulting in differential positioning of the Golgi and ERES in G1 versus S/G2 phases. In G1, the Golgi and ERES are compacted by the minus-end-directed activity of dynein and KIFC3, respectively. In S/G2, organelle-specific combinations of kinesin-1 heavy and light chains overpower minus-end-directed motors. Our findings reveal a novel, cell cycle-regulated mechanism that coordinates organelle positioning via motor switching.

## INTRODUCTION

The early secretory pathway in eukaryotic cells relies on the coordinated function of the endoplasmic reticulum (ER) and the Golgi apparatus. Secretory cargos exit the ER at specialized domains called ER exit sites (ERES), where they are packaged into COPII-coated vesicles before trafficking to the Golgi^1^. The Golgi apparatus is a central organelle in the secretory pathway, responsible for modifying, sorting, and directing newly synthesized proteins to their final destinations. The integrity and positioning of the Golgi ribbon are critical for a range of cellular functions, including migration^2,3^, protein trafficking^4,5^, cell polarity^6^, directional secretion^7^, and proper mitotic progression^8,9^. During interphase, Golgi stacks are interconnected through tubular bridges to form a continuous ribbon-like structure localized near the centrosome. The Golgi ribbon undergoes extensive fragmentation and partial fusion with the ER during mitosis to ensure equal partitioning between daughter cells^8,10–13^. Following mitotic exit, dispersed Golgi fragments and ER-to-Golgi carriers are assembled into a compact Golgi ribbon with interconnected luminal spaces in the proximity of the centrosome^14–16^.

While the mechanisms governing mitotic dynamics of the Golgi have been well characterized^8,17–19^, regulation of Golgi positioning and maintenance during interphase has received far less attention. Notably, most studies on interphase Golgi assembly and localization focus on early G1, immediately following mitosis, or in terminally differentiated G0 cells, thereby leaving a gap in our understanding of Golgi regulation during later interphase periods critical for cell growth and reproduction (S and G2).

Recent work from our lab has shown that the Golgi undergoes significant remodeling during interphase. Specifically, we observed that the compact, centrosome-associated Golgi ribbon morphology typical for G1 transitions into an extended architecture in later stages: the Golgi dissociates from the centrosome in S-phase and gradually stretches around the nuclear envelope in G2^3^. This reorganization randomizes Golgi positioning in the context of the cell polarity axis and is functionally important for the back-to-front polarity of motile cells and directional cell migration^3^. We have shown that this Golgi repositioning relies on the MT cytoskeleton^3^. However, the cell-cycle-dependent mechanism driving this process is unknown.

Based on the knowledge in the field, both MT-dependent molecular motors and the membrane carriers arriving from the ERES can influence Golgi positioning. During post-mitotic Golgi assembly, Golgi stacks are actively transported toward microtubule (MT) minus ends by the major MT-dependent motor, cytoplasmic dynein. While both centrosomal and non-centrosomal MTs assist in this transport^20,21^, the highest concentration of MT minus ends is found at the centrosome, which defines the pericentrosomal Golgi position in G1. Disruption of dynein function or its interaction with the dynactin complex leads to Golgi fragmentation and dispersal throughout the cytoplasm, mimicking the effects of MT depolymerization^22,23^. A similar phenotype is observed when the dynein adaptors are inhibited^24–26^. While dynein plays a major role in post-mitotic Golgi organization, MT plus-end-directed motor kinesin-1 heavy chain and a specific isoform of its light chain subunit, KLC1D, have been shown to localize to the Golgi^27–31^. Given the well-documented Golgi positioning at MT minus ends, the role of kinesin-1 in Golgi positioning is unclear.

In addition to the direct positioning by MT motors, Golgi positioning requires the availability of post-ER membrane carriers and is therefore influenced by the location of ERES: MT disruption results in Golgi membranes clustering near ERES^5,32^. ERES, in turn, are reciprocally sensitive to changes in Golgi organization^33,34^. In interphase mammalian cells, ERES are predominantly concentrated near the Golgi but also extend along peripheral MTs^34–36^. Similar to the Golgi, ERES positioning is highly dependent on the integrity of the MT network, as MT depolymerization acutely inhibits ERES motility^34–37^. Moreover, such motors as kinesin-1, kinesin-2, and dynein were shown to contribute to ERES dynamics^34,38^. However, to date, the mechanistic regulation of motor-dependent ERES distribution in cells under physiological conditions remains an open question. Resolving this question is even more critical, considering that assessing the role of molecular motors in Golgi positioning must account for the indirect impact of the motor-dependent location of ERES.

Based on the data outlined above, the transition of the Golgi position and morphology between the compact, polarized organization typical for G1 and extended, centrosome-detached organization^3^ in S/G2 likely depends on MT-dependent molecular motors. Moreover, it is essential to identify the molecular motors that drive the transport of both Golgi and ERES in specific substages of interphase.

To address this critical gap in knowledge, here we combined targeted loss-of-function of molecular motors with quantitative live-cell and super-resolution microscopy approaches to characterize dramatic cell cycle-dependent changes in Golgi and ERES positioning, with a particular focus on interphase substages. We demonstrate that these organelles are positioned by a dynamic tug-of-war between specific pairs of plus- and minus-end-directed MT motors (kinesin-1/dynein for the Golgi and kinesin-1/KIFC3 for ERES), modulated by a major cell cycle kinase CDK1, starting at the onset of S-phase and progressing throughout G2. Together, our findings highlight a previously unrecognized, cell cycle-regulated layer of control in the positioning of organelles.

## Results

### ERES and Golgi are differentially positioned during interphase substages

Analysis of fixed, synchronized populations of human immortalized RPE1 cells revealed distinct cell cycle-specific differences in the organization of major trafficking organelles, specifically, the Golgi complex and ERES. RPE1 cells constitutively expressing a GFP fusion of centriolar protein centrin-1^39^ were synchronized using double thymidine block^40^ and released for specific periods to analyze cells at various cell cycle substages. The substages were additionally verified by the organization of the centrosome as G1 cells contained two individual centrioles while G2 cells displayed two centriolar doublets (diplosomes), the S-phase contained either two centrioles or diplosomes with very small daughter centrioles^39,41^.

Using 3D confocal or super-resolution NSPARC imaging, we observed that the Golgi appeared compact and closely associated with the centrosome in G1 cells (Fig. 1A), consistent with previous studies^3,42^. In contrast, during the S (Supplementary Fig. 1A) and G2 (Fig. 1B) phases, the Golgi detached from the centrosomes and gradually acquired extended morphology. To quantify the spatial proximity of the centrosome to the Golgi, we calculated the mean distances from the centriole to each voxel within the segmented volumes of the Golgi (Golgi-centrosome distance, GC) (Supplementary Fig. 1B)^43^. GC values were low in the G1 phase but higher and more variable in S and G2 phase cells, indicating that the Golgi spread away from the centrosome throughout S and G2 (Fig. 1C), consistent with our prior finding^3^. Interestingly, the ERES clustered in the juxtanuclear region around the Golgi in G1 cells (Fig. 1A), while in S (Supplementary Fig. 1A) and G2 cells (Fig. 1B), they appear significantly distributed peripherally throughout the cytoplasm. To quantify the spatial proximity of the ERES to the Golgi, we measured the mean distances from the Golgi centroid to each voxel within the segmented volumes of the ERES locations (Golgi-ERES distance, GE) (Supplementary Fig. 1B). GE values were low in G1 phase but higher in S/G2 phase cells (Fig. 1D), supporting that the ERES were dispersed away from the Golgi in the later stages of the interphase.

**Figure 1.**
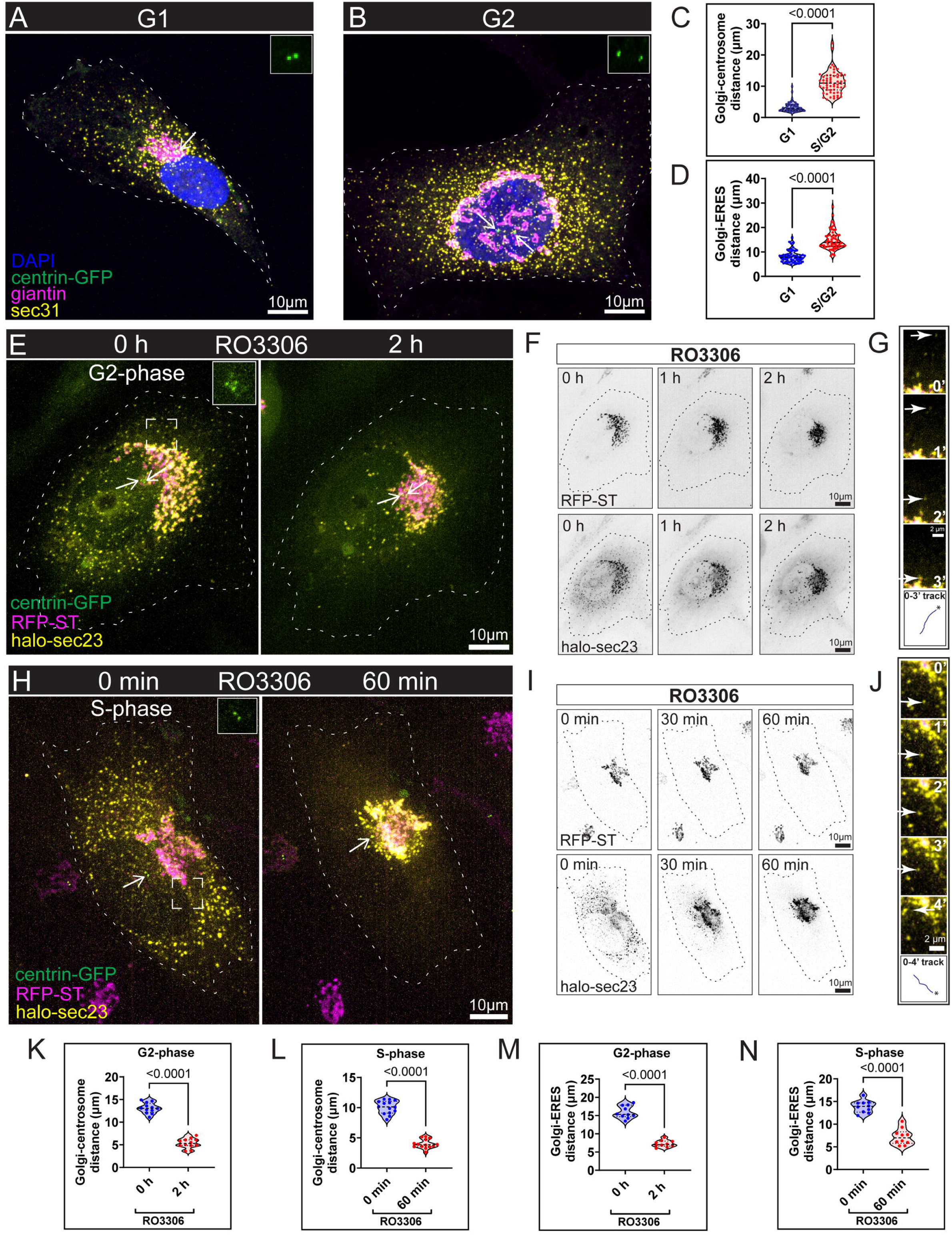
Golgi and ERES configurations are cell cycle-dependent and repositioned downstream of CDK1 in the S-phase. A, B. Representative images showing Golgi and ERES positioning in G1 (A) and G2 (B) phases. RPE1 cells stably expressing centrin-GFP (green) were immunostained for DAPI (blue), giantin (magenta), and sec31 (yellow). Images are maximum intensity projections of NSPARC stacks. Arrows indicate centrosomes, which are enlarged in solid boxed insets. Scale bar: 10 µm. C. Quantification of Golgi-centrosome distance in the fixed cell population captured on a spinning disk confocal. n=120 cells total from three independent experiments. Student’s t-test, P<0.0001. D. Quantification of Golgi-ERES distances in the fixed cell population captured on a spinning disk confocal. n=120 cells total from three independent experiments. Student’s t-test, P<0.0001. E. Time-lapse frames from live-cell imaging of RPE1 cells synchronized in the G2-phase, stably expressing Golgi (RFP-ST, magenta), centrosome (centrin-GFP, green), and ectopically expressed ERES (halo-sec23, yellow) markers treated with RO3306 (CDK1 inhibitor) for 2 h. Maximum intensity projections of spinning disk confocal stacks are shown. Arrows indicate centrosomes, which are enlarged in solid boxed insets. Scale bar: 10 µm. F. Inverted grayscale confocal images of the same cell as shown in E at indicated time points to highlight the dynamics of Golgi (RFP-ST) and ERES (halo-sec23) upon RO3306 treatment. Scale bar: 10 µm. G. Montage of the dashed boxed region in E showing retrograde ERES movement (arrows) towards the Golgi following RO3306 treatment at 1 min intervals (0-3 min). Bottom panel shows the ERES speckle track. The asterisk (*) denotes the initial position. Scale bar: 2 µm. H. Time-lapse frames from live-cell imaging of RPE1 cells synchronized in the S-phase, stably expressing Golgi (RFP-ST, magenta), centrosome (centrin-GFP, green), and ectopically expressed ERES (halo-sec23, yellow) markers treated with RO3306 (CDK1 inhibitor) for 60 min. Maximum intensity projections of spinning disk confocal stacks are shown. Arrows indicate centrosomes, which are enlarged in solid boxed insets. Scale bar: 10 µm. I. Inverted grayscale confocal images of the same cell as shown in H at indicated time points to highlight the dynamics of Golgi (RFP-ST) and ERES (halo-sec23) upon RO3306 treatment. Scale bar: 10 µm. J. Montage of the dashed boxed region in H showing retrograde ERES movement (arrows) towards the Golgi following RO3306 treatment at 1 min intervals (0-4 min). Bottom panel shows the ERES speckle track. The asterisk (*) denotes the initial position. Scale bar: 2 µm. K. Quantification of Golgi-centrosome distance in G2-phase synchronized, RO3306-treated live-cell population, based on data as shown in E. n=12 cells. Student’s t-test, P<0.0001. L. Quantification of Golgi-ERES distance in G2-phase synchronized, RO3306-treated live-cell population, based on data as shown in E. n=10 cells. Student’s t-test, P<0.0001. M. Quantification of Golgi-centrosome distance in S-phase synchronized, RO3306-treated live-cell population, based on data as shown in H. n=14 cells. Student’s t-test, P<0.0001. N. Quantification of Golgi-ERES distance in S-phase synchronized, RO3306-treated live-cell population, based on data as shown in H. n=10 cells. Student’s t-test, P<0.0001.

Overall, fixed-cell analyses showed that both the trafficking organelles, the Golgi complex and the ERES, underwent dynamic repositioning across the interphase stages.

### CDK1 activity is required to maintain the differential positioning of the Golgi and ERES in S-phase

CDK1, a serine/threonine kinase, serves as a master regulator of the cell cycle^44–47^. Its activity begins to rise during the S phase, peaks in metaphase, and declines rapidly in anaphase^48–50^. Because this increase in activity coincides with the progressive Golgi/ERES redistribution, we investigated whether CDK1 regulates organelle positioning. We performed 3D live-cell time-lapse imaging of RPE1 cells co-expressing centrin1-GFP (centrosome marker), RFP-ST (Golgi marker), and Halo-sec23 as an ERES marker to monitor the positioning of Golgi and ERES. In G2-synchronized cells, acute inhibition of CDK1 activity using RO3306, a selective inhibitor of CDK1^51^, induced retrograde repositioning of both the Golgi and ERES, leading to a G1-like compact, pericentrosomal Golgi/ERES organization (Supplementary video 1, Fig 1E-G). Strikingly, acute CDK1 inhibition in S-phase facilitated similar compaction of Golgi/ERES (Fig. 1H-J and Supplementary video 2), indicating that modestly rising levels of CDK1 activity in S-phase are sufficient to trigger the morphological transition. A drastic reduction in both the GC (Fig. 1K-L) and GE (Fig. 1M-N) distances upon RO3306 treatment corroborated this morphological shift, highlighting the role of CDK1 in organelle positioning in the S-phase and G2.

### Kinesin-1 positions Golgi and ERES in the S/G2 phase

MT-dependent motors play a crucial role in the spatial arrangement of membrane-bound organelles^52–54^. Thus, we hypothesized that motors may selectively promote cell-cycle-dependent repositioning of the Golgi and ERES. Because the centrifugal redistribution of organelles during interphase transitions suggests a role of a plus-end directed motor, and kinesin-1 was previously implicated in Golgi morphology and ERES transport^27–29,34,38^, we tested kinesin-1 involvement in organelle positioning in S/G2 cells. We disrupted the cargo-binding function of kinesin-1 using kinesore, a small molecule modulator of kinesin-1 function^55–57^. Such acute inhibition of kinesin-1-driven transport induced active retrograde movement of both the Golgi and ERES, mirroring the effects of CDK1 inhibition (Fig. 2A-B and Supplementary video 3). Specifically, the equatorially distributed Golgi compacted and clustered around the centrosomes, while the peripheral ERES relocalized to perinuclear Golgi regions (Supplementary video 3). Accordingly, both the GC (Fig. 2C) and GE (Fig. 2D) distances were significantly reduced after kinesore treatment. The directional movement of individual ERES puncta towards the Golgi cluster indicated the active transport nature of this reorganization (Fig. 2B).

**Figure 2.**
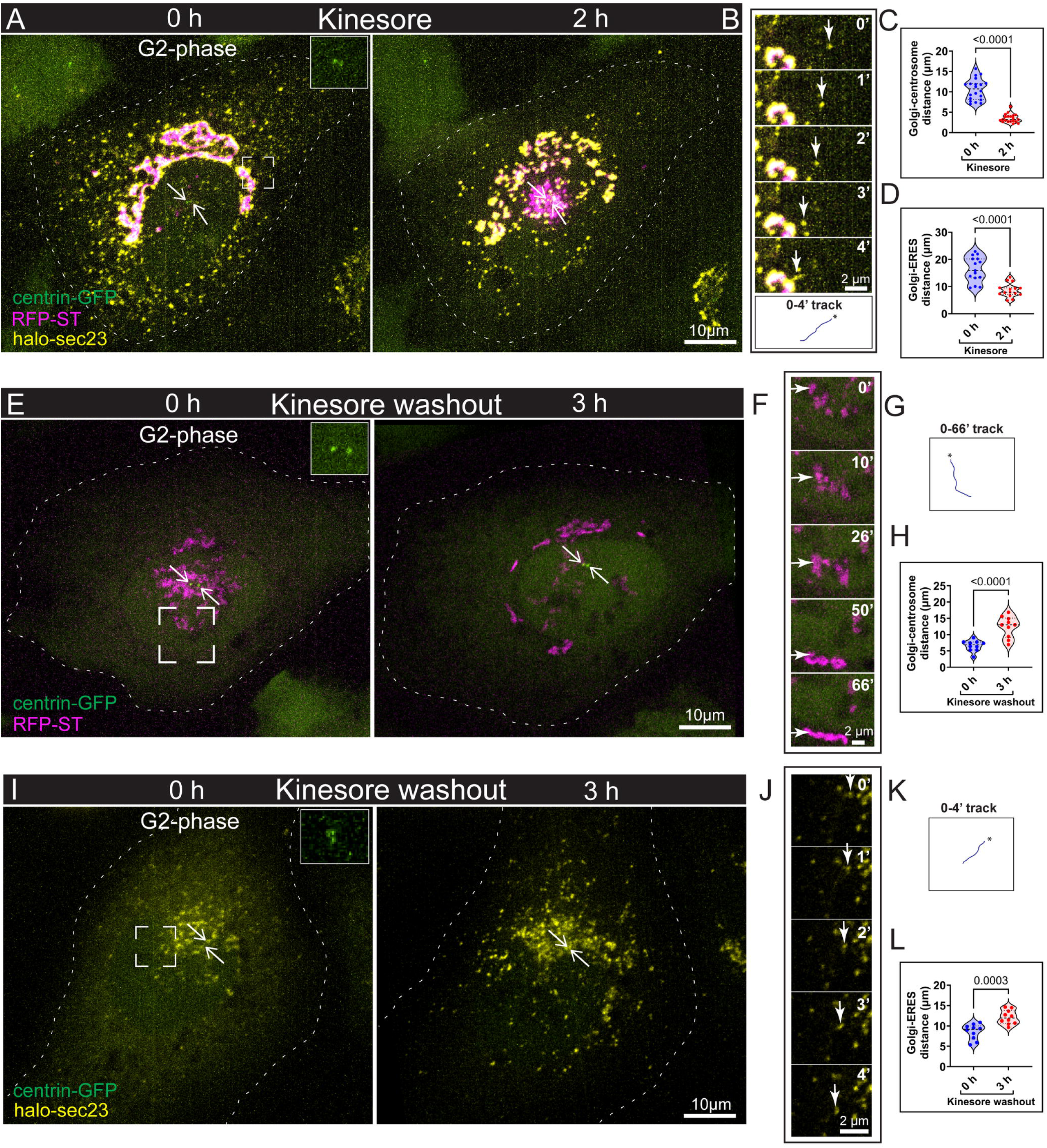
Kinesin-1 positions Golgi and ERES in the S/G2 phase. A. Time-lapse frames from live-cell imaging of RPE1 cells synchronized in the G2-phase, stably expressing Golgi (RFP-ST, magenta), centrosome (centrin-GFP, green), and ectopically expressed ERES (halo-sec23, yellow) markers treated with kinesore (kinesin-1 modulator) for 2 h. Maximum intensity projections of spinning disk confocal stacks are shown. Arrows indicate centrosomes, which are enlarged in solid boxed insets. Scale bar: 10 µm. B. Montage of the dashed boxed region shown in A, illustrating retrograde movement of ERES (arrows) towards the Golgi upon kinesore treatment at 1 min intervals (0-4 min). 0 to 4 min track of ERES speckle on kinesore treatment is shown in the bottom panel. The asterisk (*) denotes the initial position. Scale bar: 2 µm. C. Quantification of Golgi-centrosome distance in S- and G2-phase synchronized, kinesore-treated live-cell population, based on data as shown in A. n=20 cells. Student’s t-test, P<0.0001. D. Quantification of Golgi-ERES distance in S- and G2-phase synchronized, kinesore-treated live-cell population, based on data as shown in A. n=15 cells. Student’s t-test, P<0.0001. E. Time-lapse frames from live-cell imaging of RPE1 cells synchronized in the G2-phase, stably expressing Golgi (RFP-ST, magenta) and centrosome (centrin-GFP, green) markers treated with kinesore for 2 h, washed, and incubated in kinesore-free medium for 3 h. Maximum intensity projections of spinning disk confocal stacks are shown. Arrows indicate centrosomes, which are enlarged in solid boxed insets. Scale bar: 10 µm. F. Montage of the dashed boxed region shown in E, illustrating anterograde movement of Golgi stacks (arrows) away from centrosomes upon kinesore washout over time. Scale bar: 2 µm. G. 0 to 66 min track of Golgi stack (shown in Fig. F) movement on kinesore washout is shown. The asterisk (*) denotes the initial position. H. Quantification of Golgi-centrosome distance in S- and G2-phase synchronized, kinesore-washout live-cell population, based on data as shown in E. n= 10 cells. Student’s t-test, P<0.0001. I. Time-lapse frames from live-cell imaging of RPE1 cells synchronized in the G2-phase, stably expressing centrosome (centrin-GFP, green) and ectopically expressed ERES (halo-sec23, yellow) markers treated with kinesore for 2 h, washed, and incubated in kinesore-free medium for 3 h. Maximum intensity projections of spinning disk confocal stacks are shown. Arrows indicate centrosomes, which are enlarged in solid boxed insets. Scale bar: 10 µm. J. Montage of the dashed boxed region shown in I, illustrating anterograde movement of ERES (arrows) away from centrosomes upon kinesore washout at 1 min intervals (0-4 min). Scale bar: 2 µm. K. 0 to 4 min track of ERES speckle (shown in Fig. J) on kinesore washout is shown. The asterisk (*) denotes the initial position. L. Quantification of Golgi-ERES distance in S- and G2-phase synchronized, kinesore-washout live-cell population, based on data as shown in I. n= 10 cells. Student’s t-test, P<0.001.

Strikingly, these effects were reversible: kinesore washout in G2 resulted in the anterograde movement of the Golgi, detaching it from the centrosomes and redistributing around the nuclear envelope (Fig. 2E-G and Supplementary video 4). Similarly, ERES underwent anterograde movement from their accumulations near the Golgi and redistributed throughout the cytoplasm (Fig. 2I-K and Supplementary video 5). Correspondingly, GC (Fig. 2H) and GE distances (Fig. 2L) also returned to S/G2-like values. In addition, kinesore washout enhanced the sec23 signal at ERES, indicating a positive role of kinesin-1 in ERES function (Supplementary video 5). These findings demonstrated that kinesin-1 is essential for maintaining the spatial organization of both the Golgi and ERES during the S/G2 phase of the cell cycle.

### KIF5B positions the Golgi but not ERES during the S/G2 phase

Each kinesin-1 molecule includes two heavy (motor) chains and two light chains for cargo attachment. In mammalian cells, there are three KHCs (KIF5A, KIF5B, and KIF5C) and four KLC (KLC1–4) variants encoded by distinct genes^58^. Thus, we proceeded to identify which kinesin-1 variants are responsible for positioning the Golgi and ERES during the S/G2 phase.

To directly test if organelle positioning relies on KIF5B, which out of three KHCs has the highest expression levels in RPE1 cells, we used a CRISPR-mediated KIF5B knockout (KIF5B KO) cell line^59^. Our analyses revealed that in the absence of KIF5B, the Golgi in G2/S cells adopted a compact configuration clustered at the centrosome, indicating that KIF5B is critical for maintaining an equatorial distribution of Golgi during these phases (Fig. 3A-C). In contrast, ERES largely retained their peripheral distribution, suggesting that KIF5B was dispensable for ERES positioning (Fig. 3A, B, D). Furthermore, in KIF-5B KO cells in G2, treatments with both CDK1 inhibitor RO3306 and kinesin-1 transport inhibitor kinesore had no effect on the already clustered Golgi, while ERES underwent retrograde repositioning toward the perinuclear region (Fig. 3E-J and Supplementary videos 6 and 7). These findings demonstrated that KIF5B is essential for the Golgi extension in the S/G2 phase. In contrast, the continued sensitivity of ERES to RO3306 and kinesore in KIF5B KO cells suggested that ERES positioning relies on a different kinesin-1 variant.

**Figure 3.**
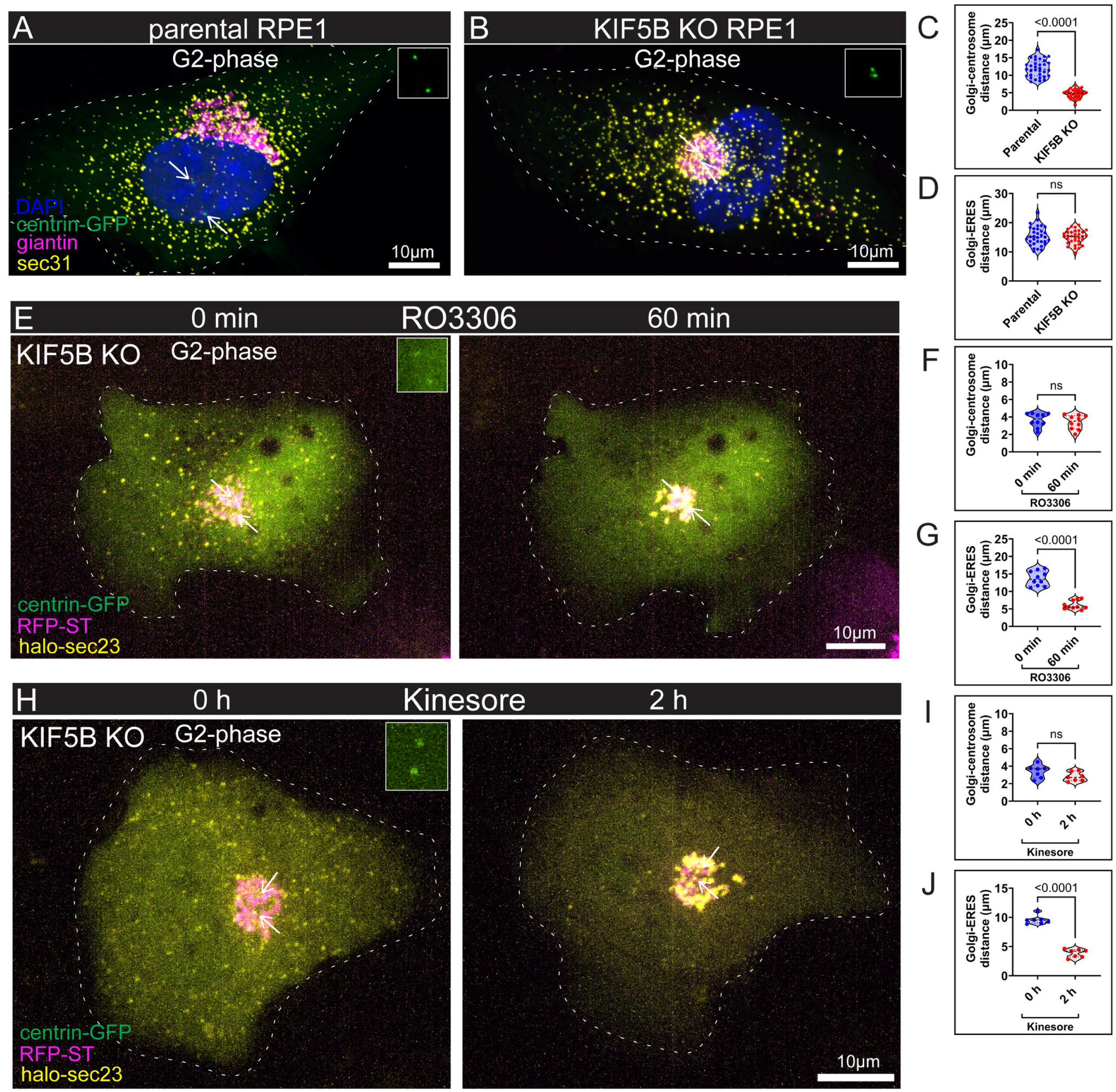
KIF5B positions the Golgi but not ERES during the S/G2 phase. A, B. Representative images showing Golgi and ERES positioning in parental (A) and KIF5B KO RPE1 (B) cells synchronized in the G2 phase. Parental and KIF5B KO RPE1 cells expressing centrin-GFP (green) were immunostained for DAPI (blue), giantin (magenta), and sec31 (yellow). Images are maximum intensity projections of laser scanning confocal stacks. Arrows indicate centrosomes, which are enlarged in solid boxed insets. Scale bar: 10 µm. C. Quantification of Golgi-centrosome distance in S- and G2-phase synchronized, fixed cell population, based on data as shown in A and B. n=60 cells total from three independent experiments. Student’s t-test, P<0.0001.D. Quantification of Golgi-ERES distance in S- and G2-phase synchronized, fixed cell population, based on data as shown in A and B. n=60 cells total from three independent experiments. Student’s t-test, ns: P>0.05. E. Time-lapse frames from live-cell imaging of KIF5B KO RPE1 cells synchronized in the G2-phase, ectopically expressing Golgi (RFP-ST, magenta), centrosome (centrin-GFP, green), and ERES (halo-sec23, yellow) markers treated with RO3306 for 60 min. Maximum intensity projections of spinning disk confocal stacks are shown. Arrows indicate centrosomes, which are enlarged in solid boxed insets. Scale bar: 10 µm. F. Quantification of Golgi-centrosome distance in S- and G2-phase synchronized, RO3306-treated live-cell population, based on data as shown in E. n=10 cells. Student’s t-test, ns: P>0.05. G. Quantification of Golgi-ERES distance in S and G2-phase synchronized, RO3306-treated live-cell population, based on data as shown in E. n=10 cells. Student’s t-test, P<0.0001. H. Time-lapse frames from live-cell imaging of KIF5B KO RPE1 cells synchronized in the G2-phase, ectopically expressing Golgi (RFP-ST, magenta), centrosome (centrin-GFP, green), and ERES (halo-sec23, yellow) markers treated with kinesore for 2 h. Maximum intensity projections of spinning disk confocal stacks are shown. Arrows indicate centrosomes, which are enlarged in solid boxed insets. Scale bar: 10 µm. I. Quantification of Golgi-centrosome distance in S- and G2-phase synchronized, kinesore-treated live-cell population, based on data as shown in H. n=7 cells. Student’s t-test, ns: P>0.05. J. Quantification of Golgi-ERES distance in S and G2-phase synchronized, kinesore-treated live-cell population, based on data as shown in H. n=7 cells. Student’s t-test, P<0.0001.

### KLC1 is essential for Golgi organization and KLC3 for ERES organization in S/G2 cells

To further dissect the motor complex involved, we next examined the role of specific KLC isoforms in the spatial organization of these trafficking organelles. Depletion of KLC1 (Supplementary Fig. 2A and B) during the S/G2 phase resulted in a centrally collapsed Golgi tightly clustered around the centrosomes (Fig. 4A-C). In contrast, ERES distribution remained largely unaltered upon KLC1 knockdown (Fig. 4A, B, D). This phenotype was reminiscent of the phenotype observed in KIF5B KO cells (Fig. 3A-D), while KIF5B protein levels remained unchanged (Supplementary Fig. 2C and D). To determine whether other kinesin light chains contributed to ERES positioning, we next depleted KLC3 (Supplementary Fig. 2E and F). Fixed-cell imaging showed a marked perinuclear accumulation of ERES near the Golgi, indicating that KLC3 is required for maintaining peripheral ERES distribution (Fig. 4E-F and H). Intriguingly, KLC3 knockdown also resulted in Golgi compaction (Fig. 4F-G). Further immunoblot analysis revealed a reduction in KLC1 protein levels upon KLC3 depletion (Supplementary Fig. 2G and H), raising the possibility of regulatory crosstalk between light chain isoforms, a phenomenon that warranted further investigation. Together, these data indicated that KLC1 is essential for KIF5B-dependent positioning of the Golgi, while KLC3 is needed for kinesin-1-dependent ERES positioning.

**Figure 4.**
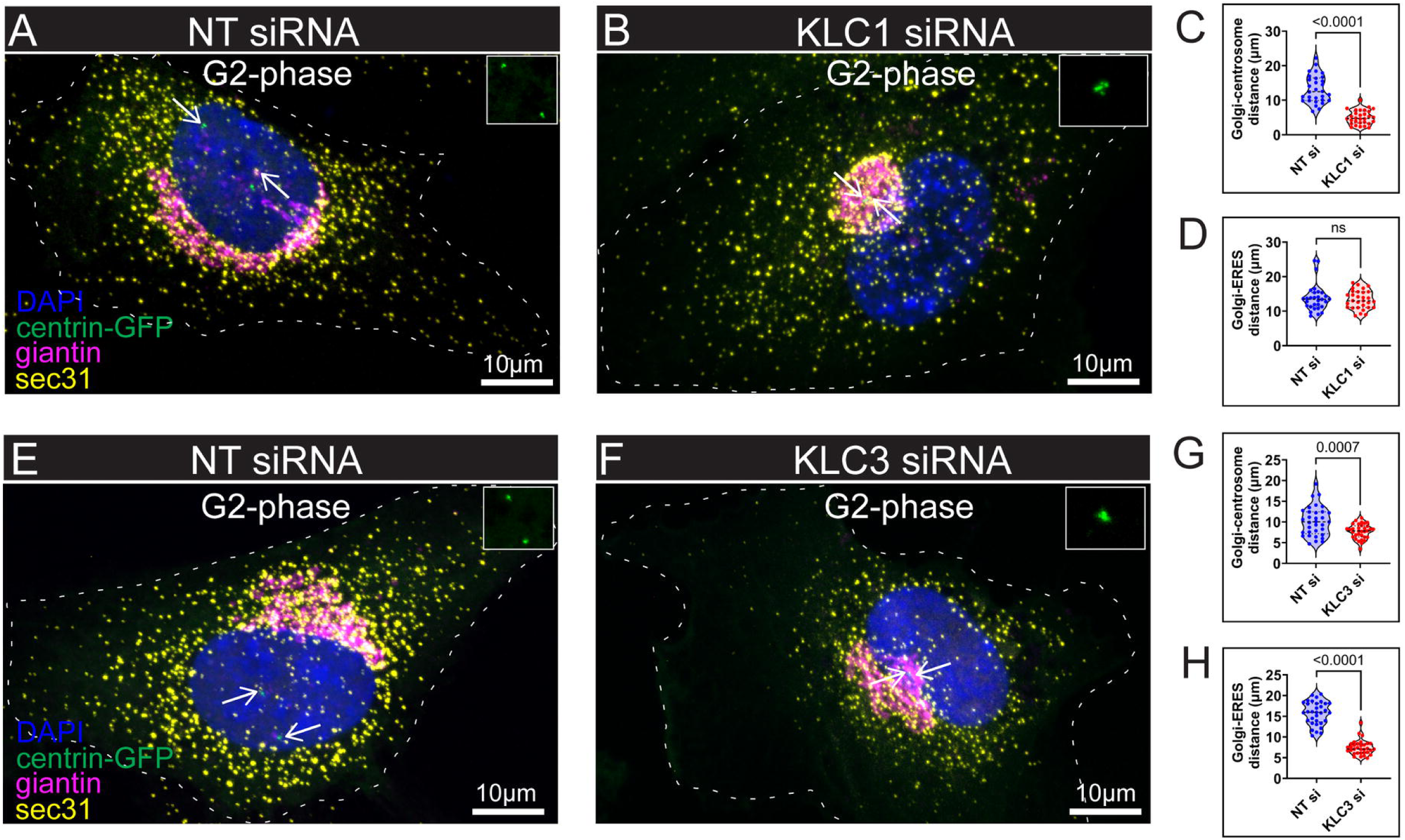
KLC1 is essential for Golgi organization and KLC3 for ERES organization in S/G2 cells. A, B. Representative images of Golgi and ERES positioning in non-targeting (NT) control (A) and KLC1 siRNA-treated (B) RPE1 cells synchronized in G2 phase. RPE1 cells stably expressing centrin-GFP (green) were immunostained for DAPI (blue), giantin (magenta), and sec31 (yellow). Images are maximum intensity projections of laser scanning confocal stacks. Arrows indicate centrosomes, which are enlarged in solid boxed insets. Scale bar: 10 µm. C. Quantification of Golgi-centrosome distance in S- and G2-phase synchronized, fixed cell population, based on data as shown in A and B. n=60 cells total from three independent experiments. Student’s t-test, P<0.0001. D. Quantification of Golgi-centrosome and Golgi-ERES distances in S- and G2-phase synchronized, fixed cell population, based on data as shown in A and B. n=60 cells total from three independent experiments. Student’s t-test, ns: P>0.05. E, F. Representative images of Golgi and ERES positioning in NT control (E) and KLC3 siRNA-treated (F) RPE1 cells synchronized in G2 phase. RPE1 cells stably expressing centrin-GFP (green) were immunostained for DAPI (blue), giantin (magenta), and sec31 (yellow). Images are maximum intensity projections of laser scanning confocal stacks. Arrows indicate centrosomes, which are enlarged in solid boxed insets. Scale bar: 10 µm. G. Quantification of Golgi-centrosome distance in S- and G2-phase synchronized, fixed cell population, based on data as shown in E and F. n=60 cells total from three independent experiments. Student’s t-test, P<0.001. H. Quantification of Golgi-ERES distance in S- and G2-phase synchronized, fixed cell population, based on data as shown in E and F. n=60 cells total from three independent experiments. Student’s t-test, P<0.0001.

### Kinesin-1 is counterbalanced by dynein for Golgi positioning and by KIFC3 for ERES positioning

The retrograde repositioning of both Golgi and ERES upon inactivating kinesin-1 in G2 suggested that minus-end-directed motors at these organelles remain active at this cell-cycle phase. Previous studies have implicated dynein and its regulatory proteins in Golgi membrane trafficking and maintenance of Golgi ribbon structure and positioning^22,23^. Thus, we assessed whether dynein actively contributed to Golgi and ERES positioning during the S/G2 phase and whether it engaged in a possible tug-of-war with kinesin-1. We treated S/G2 cells with RO3306 to induce the reversal of organelle positioning to the G1-like state, at the same time suppressing dynein-mediated retrograde transport by ciliobrevin D, a dynein ATPase inhibitor^60,61^. Upon dual treatment, we observed pronounced retrograde movement of ERES towards perinuclear Golgi regions, similar to the phenotype seen with CDK1 inhibition alone (Fig. 5A and supplementary video 8). The Golgi, however, remained immobile, extended, and detached from the centrosome but appeared fragmented (Fig. 5A and supplementary video 8). Consistent with this, GC distances remained largely unchanged (Fig. 5B), while GE values significantly decreased (Fig. 5C). To further dissect the contributions of dynein and kinesin-1 to Golgi positioning, we treated cells with both ciliobrevin D and kinesore. This dual inhibition resulted in similar fragmented and immobile Golgi localized near the nuclear envelope as above, while peripheral ERES exhibited retrograde movement towards the Golgi (Fig. 5D-F and supplementary video 9). These findings support a model in which both kinesin-1 and dynein are critical for maintaining proper Golgi positioning in S/G2 phase cells.

**Figure 5.**
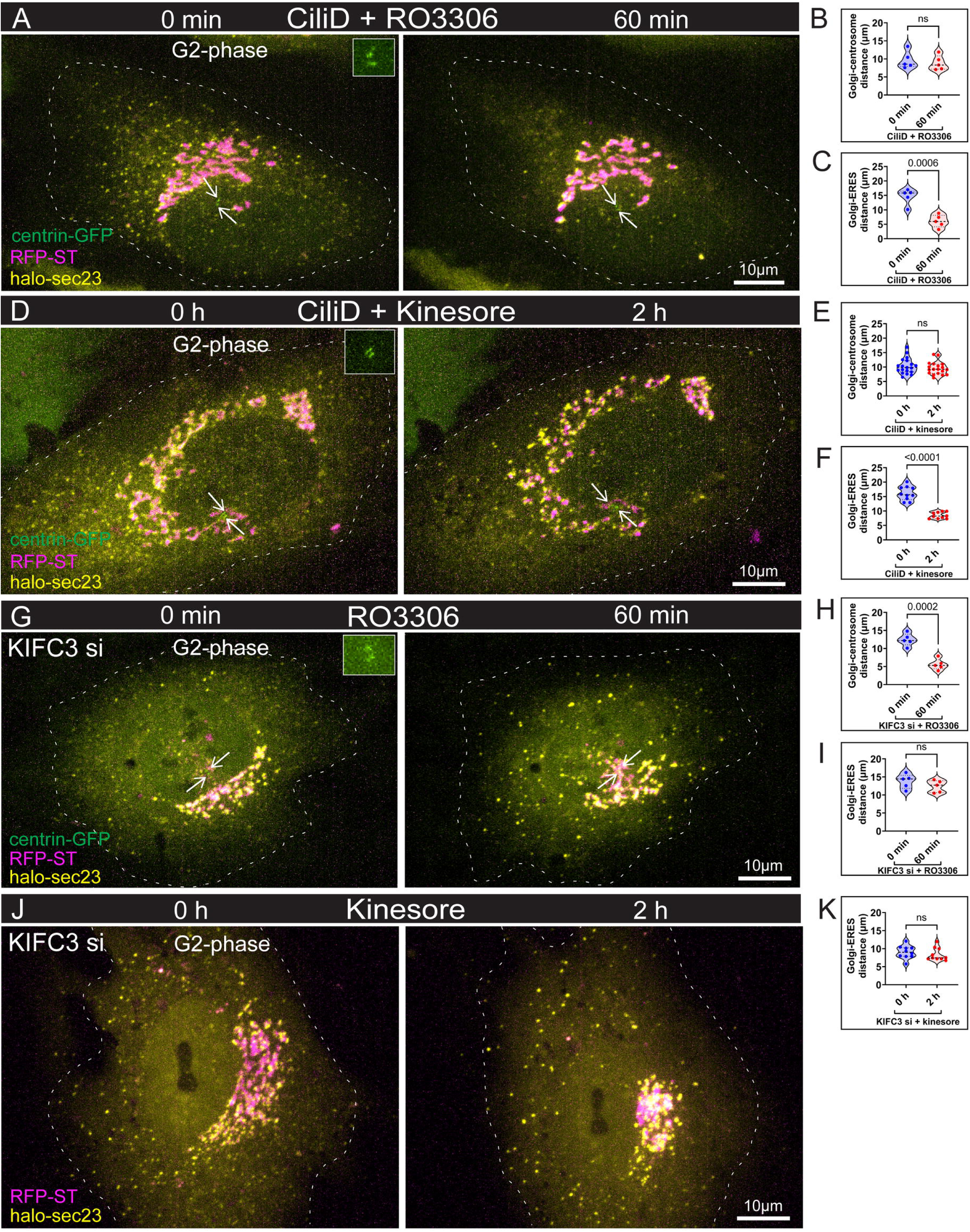
Kinesin-1 is counterbalanced by dynein for Golgi positioning and by KIFC3 for ERES positioning. A. Time-lapse frames from live-cell imaging of RPE1 cells synchronized in the G2-phase, stably expressing Golgi (RFP-ST, magenta), centrosome (centrin-GFP, green), and ectopically expressed ERES (halo-sec23, yellow) markers treated with Ciliobrevin D (CiliD, dynein inhibitor) and RO3306 for 1 h. Maximum intensity projections of spinning disk confocal stacks are shown. Arrows indicate centrosomes, which are enlarged in solid boxed insets. Scale bar: 10 µm. B. Quantification of Golgi-centrosome distance in S- and G2-phase synchronized, CiliD and RO3306-treated live-cell population, based on data as shown in A. n=5 cells. Student’s t-test, ns: P>0.05. C. Quantification of Golgi-ERES distance in S- and G2-phase synchronized, CiliD and RO3306-treated live-cell population, based on data as shown in A. n=5 cells. Student’s t-test, P<0.001. D. Time-lapse frames from live-cell imaging of RPE1 cells synchronized in the G2-phase, stably expressing Golgi (RFP-ST, magenta), centrosome (centrin-GFP, green), and ectopically expressed ERES (halo-sec23, yellow) markers treated with CiliD and kinesore for 2 h. Maximum intensity projections of spinning disk confocal stacks are shown. Arrows indicate centrosomes, which are enlarged in solid boxed insets. Scale bar: 10 µm. E. Quantification of Golgi-centrosome distance in S- and G2-phase synchronized, CiliD and kinesore-treated live-cell population, based on data as shown in D. n=19 cells. Student’s t-test, ns: P>0.05. F. Quantification of Golgi-ERES distances in S- and G2-phase synchronized, CiliD and kinesore-treated live-cell population, based on data as shown in D. n=10 cells. Student’s t-test, P<0.0001. G. Time-lapse frames from live-cell imaging of KIFC3 knockdown RPE1 cells synchronized in the G2-phase, stably expressing Golgi (RFP-ST, magenta), centrosome (centrin-GFP, green), and ectopically expressed ERES (halo-sec23, yellow) markers treated with RO3306 for 1 h. Maximum intensity projections of spinning disk confocal stacks are shown. Arrows indicate centrosomes, which are enlarged in solid boxed insets. Scale bar: 10 µm. H. Quantification of Golgi-centrosome in S- and G2-phase synchronized, RO3306-treated KIFC3 knockdown live-cell population, based on data as shown in G. n=5 cells. Student’s t-test, P<0.001. I. Quantification of Golgi-ERES distance in S- and G2-phase synchronized, RO3306-treated KIFC3 knockdown live-cell population, based on data as shown in G. n=5 cells. Student’s t-test, ns: P>0.05. J. Time-lapse frames from live-cell imaging of KIFC3 knockdown RPE1 cells synchronized in the G2-phase, stably expressing Golgi (RFP-ST, magenta) and ectopically expressed ERES (halo-sec23, yellow) markers treated with kinesore for 2 h. Maximum intensity projections of spinning disk confocal stacks are shown. Scale bar: 10 µm. K. Quantification of Golgi-ERES distance in S- and G2-phase synchronized, kinesore-treated KIFC3 knockdown live-cell population, based on data as shown in J. n=10 cells. Student’s t-test, ns: P>0.05.

To identify the motor complex responsible for ERES retrograde transport, we next examined the roles of minus-end-directed kinesins. KIFC3-depleted S/G2 cells (Supplementary Fig. 2I and J) were treated with RO3306 to induce retrograde organelle repositioning. Under these conditions, the Golgi repositioned and clustered around centrosomes, mirroring phenotype upon CDK1 inhibition; however, the retrograde movement of ERES markedly reduced (Fig. 5G-I and supplementary video 10). To further explore the interplay between KIFC3 and kinesin-1, we treated KIFC3-depleted cells with kinesore. As expected, we observed Golgi compaction, consistent with kinesore treatment; however, retrograde repositioning of ERES was reduced (Fig. 5J-K and supplementary video 11), suggesting a tug-of-war between KIFC3 and kinesin-1 in regulating ERES positioning in cell cycle phases.

Together, these findings suggested that while dynein and kinesin-1 exert opposing forces to regulate Golgi positioning in S/G2, ERES relies on a distinct motor pair, kinesin-1 and KIFC3, for its spatial distribution. These results underscored that the differential positioning of the Golgi and ERES in S/G2 was mediated by their selective engagement with distinct sets of molecular motors.

## DISCUSSION

In this study, we highlight the dynamic and cell cycle-regulated organization of the Golgi apparatus and ERES and identify the molecular players and mechanisms that control their transitions. We demonstrate that distinct MT-dependent molecular motors promote a centralized pericentrosomal position of both organelles in G1 and an extended distribution in S-phase and G2. Specifically, Golgi positioning is driven by kinesin-1 and dynein, while ERES positioning is driven by kinesin-1 and KIFC3.

Both the redistribution of ERES throughout the cytoplasm and the detachment of the Golgi from the centrosome, observed at the onset of S-phase, can be reversed by the kinesin-1 detachment from the membranes. This indicates that the transition between compact and extended organelle configurations is potentiated by kinesin activity at the Golgi and ERES, overpowering minus-end-directed motors. This supports a striking model of motor coordination via a tug-of-war mechanism, wherein kinesin-1 and minus-end-directed motors simultaneously bind to membranes and exert opposing forces. Prior *in vitro* and *in vivo* studies have shown that antagonistic motors can co-exist on the same cargo^62–65^, with bidirectional transport regulated by their coordination. The widely accepted “tug-of-war” model proposes that cargo directionality is determined by which motor force dominates^66,67^. Notable support for this model includes observations of bidirectional switches in Dictyostelium endosomes^62^ and vesicular transport in neurons^63^. Cellular studies have also provided information about the number of motors required for bidirectional motility^68,69^. Our findings provide one of the first examples of cell cycle-regulated, organelle-scale tug-of-war in a physiological context.

Unexpectedly, we find that while Golgi and ERES undergo similar transitions in the cell cycle, the movement of each organelle requires specific motors. While the minus-end-directed component of the Golgi position is modulated by dynein as established previously^22,23^, ERES are moved by the minus-end-directed kinesin KIFC3. This is a novel major role for this understudied member of the kinesin-14 family with very few reported functions (e.g. apical vesicle trafficking in polarized epithelial cells^70^ and peroxisomal transport in kidney cells^71^). Interestingly, prior studies have implicated KIFC3 in Golgi positioning in adrenocortical cells under cholesterol-depleted conditions^72^. One of the described consequences of cholesterol depletion is inefficient dynein recruitment to the membrane^73,74^. Together with our new finding, the study by Xu and Hirokawa^72^, might indicate that upon cholesterol depletion, the Golgi cannot be transported by dynein and its positioning heavily relies on the ERES location.

Furthermore, the plus-end-dependent component of both organelle positioning is driven by kinesin-1, but the motor isoforms are distinct. In agreement with previous findings, KIF5B/KLC1 is responsible for the Golgi^27–30^, but ERES positioning responds to both small-molecule inactivation of all kinesin-1-dependent trafficking and KLC3 depletion. We interpreted the sensitivity of ERES to kinesore in the absence of KIF5B as potential ERES transport by other kinesin-1 variants, KIF5A and/or KIF5C, both of which are steadily expressed in RPE1 cells, albeit at lower levels than KIF5B, as reported by Human Protein Atlas. The organelle-specific utilization of KLCs is consistent with the current understanding of KLC roles in cargo selection by kinesin-1: cargo adaptors can either selectively recruit motors (“selective recruitment”) or modulate their engagement with microtubules (“selective activation”)^75^. Engagement of different heavy chains is more puzzling and might provide a route to answer an important open question that remains to be explored: why are Golgi and ERES positioned differentially? One possibility is that the motors that move their respective organelles select different preferred MT tracks. It is known that specific motors select their preferred tracks according to the tubulin code (tubulin post-translational modifications (PTMs)^76–79^ and MT-associated proteins (MAPs)^80–83^. It is possible that the Golgi and ERES may be transported along MT subsets, specialized via these mechanisms. Interestingly, the difference between Golgi and ERES positioning is most prominent in S/G2, when kinesin-1 takes over, suggesting that different kinesin-1 isoforms might select different tracks. This is provocative because such differential track selection has been described for different kinesin families and dynein^79,84,85^ but not yet for closely related kinesins. Overall, the specialization of motors at these two interdependent organelles suggests an elegant layer of their positioning regulation at the level of motor-cargo binding.

Interestingly, we find that the balance of motors is tuned downstream of CDK1. This is one of the CDK1 functions that is already manifest in the S-phase when CDK1 activity begins to rise^48,49,86^. Several mitotic kinesins, such as Eg5^87^, MCAK^88^, and KIF18A^89^, are known to be activated by CDK1. Our finding suggests that interphase motor interplay is very sensitive to CDK1 regulation. Specific CDK1-dependent pathway controlling tug-of-war at organelles is yet to be elucidated. Most likely, this regulation targets the common denominator in Golgi and ERES positioning, specifically kinesin-1. Kinesin-1 is known to undergo auto-inhibition via folding^90–92^, and CDK1 may relieve this inhibition through direct heavy-chain phosphorylation. Such kinesin-1 activation could bias transport directionality, tilting the tug-of-war. Alternatively, CDK1 might fine-tune the balance between opposing motors by modulating their membrane association. Post-translational modifications of adaptors like phosphorylation (e.g. JIP1^93^) or the presence of binding partners (e.g. HAP1^94,95^) can modulate whether kinesin or dynein binds a given cargo. In our case, CDK-1 could enhance kinesin-1 recruitment to membranes by phosphorylating KLCs, thereby biasing motor activity toward the plus-end. It is also possible that CDK1 regulates activity or organelle attachment of minus-end directed motors: dynein was shown to be regulated by CDK1 in neurons and Xenopus extracts^96–99^.

Another important consideration is how the organelle positioning affects their roles in cell physiology. We and others have previously shown that dissociation of the Golgi from the centrosomes is essential for centrosome separation in preparation for mitosis^3,19^. Also, we speculate that the change in the mutual positioning of the Golgi and ERES has a function in membrane trafficking. Shortening the distance between ERES and the Golgi has been proposed to enhance cargo delivery^34,100^. However, we observe that in S/G2, ERES become more scattered throughout the cytoplasm. This might indicate that rapid ER-to-Golgi transport is more important in G1 than during preparation for mitosis. Alternatively, the difference in ERES pattern could reflect a different range of cargos required in different cell cycle phases: ERES are known to show adaptability to diverse cargo sizes and export rates^100,101^, and a more dispersed ERES network could accommodate a broader distribution of transport carriers in S/G2. Also, this reorganization could favor certain cargos, which are preferentially transported through tubule-mediated rather than through short-distance ER-to-Golgi transfer^101,102^.

In sum, our study reveals how motor-dependent cell cycle cues and motor coordination govern Golgi and ERES reorganization. We propose that a CDK1-regulated switch from minus- to plus-end motor dominance underlies these transitions and that specific kinesin-1 light chain isoforms act as critical mediators of compartment-specific positioning. These findings open new avenues for exploring how the spatial organization of organelles contributes to essential cellular functions, especially in contexts of secretion, cell migration, and division.

## Materials and Methods

### Cell Culture

All cell lines were maintained in a humidified incubator at 37 °C in 5% CO_2_. Immortalized human retinal pigment epithelial hTERT-RPE1 cells (CRL-4000, ATCC) were grown in DMEM/F12 media (Life Technologies, Frederick, MD) with 10% Fetal Bovine Serum (FBS), 100 µM penicillin, and 0.1 mg/mL streptomycin (Fisher Scientific, Waltham, MA). KIF5B knockout (KO) RPE1 cells were obtained from Vladimir Gelfand and maintained under the same conditions^59^. Either of the two hTERT-RPE1 stable cell lines was used for most experiments. (1) Centrin-1-GFP was expressed by lentiviral incorporation of the construct^39^ (gifted from Alexey Khodjakov) into the wild-type RPE1 cells (2) Centrin-1-GFP RPE1 cells were then transfected with RFP-ST^103^ (gifted from Enrique Rodriguez-Boulan), via Amaxa (Lonza, Walkersville, MD), then selected with G418 for at least two weeks before cell sorting.

### Synchronization of RPE1 cells by double thymidine block

To synchronize the RPE1 cell population, a standard double thymidine block protocol was employed^40^. Briefly, cells were incubated in a culture medium supplemented with 2 mM thymidine (#T1895, Sigma-Aldrich, St. Louis, MO) for 24 h. After an initial block, cells were released into a thymidine-free culture medium for 8 h, followed by a second incubation in 2 mM thymidine for 18 h. Finally, cells were released into a fresh medium and collected at different time points to enrich for specific cell cycle phases: 2 h post-release for the S phase, 6 h for the G2 phase, and 14 h for the G1 phase.

### Plasmids, siRNA constructs, and reagents

The following constructs were used: Halo-sec23a^101^ (#166892, Addgene), RFP-ST^103^ (gifted from Enrique Rodriguez-Boulan), and Centrin-1-GFP^39^ (gifted from Alexey Khodjakov).

A pool of four ON-TARGETplus siRNA sequences (SMARTPool; Dharmacon, Lafayette, CO) was used against human KLC1 (cat. no: L-019482-00-0005), human KLC3 (cat. no: L-016063-00-0005), human KIFC3 (cat. no: L-008338-00-0005). An ON-TARGETplus non-targeting control pool (Cat. No. D-001810-10-05) was used as a negative control. All plasmid and siRNA constructs were transfected using TransIT-X2 (Mirus, Madison, WI) according to the manufacturer’s protocols.

Small molecule reagents include dimethyl sulfoxide (DMSO; #T7402, Sigma-Aldrich, St. Louis, MO), Kinesore (Tocris Bioscience, Minneapolis, MN), Ciliobrevin D (#250401, Sigma-Aldrich, St. Louis, MO), RO3306 (#SML0569, Sigma-Aldrich, St. Louis, MO).

### Immunofluorescence staining

Cells were fixed in 4% paraformaldehyde, permeabilized with 0.1% Triton X-100, and later blocked in 1% DHS/BSA (Donor Horse Serum, Bio-Techne, MN; Bovine Serum Albumin, Fisher Scientific, Waltham, MA) for 1 h at room temperature. Primary antibodies included: anti-giantin (1:300, #ab80864; Abcam, Cambridge, MA), anti-sec31 (1:300, #612351; BD Biosciences, Franklin Lake, NJ). Alexa Fluor 568-, and 647-conjugated highly cross-absorbed secondary antibodies (1:500, Invitrogen, Waltham, MA) were used. DNA was stained with either DAPI (Fisher Scientific, Waltham, MA) or Hoechst 33457 (Fisher Scientific, Waltham, MA). Coverslips were mounted in Vectashield Antifade Mounting Medium (Vector Labs, Burlingame, CA).

### Image Acquisition of Fixed and Live Samples

Fixed samples were imaged using a Nikon A1R laser scanning confocal microscope equipped with a TiE motorized inverted base and an SR Apo TIRF 100X NA 1.49 lens, controlled via NIS Elements C software (Nikon, Tokyo, Japan). Z-stacks were acquired at 0.09 μm intervals, and maximum intensity projections were generated for whole-cell visualization. Super-resolution images were captured using a Nikon AX R microscope with an NSPARC detector system and a 100X NA 1.49 lens. For all live-cell imaging, cells were maintained on the microscope stage at 37 °C in 5% CO_2_. Cells were seeded on glass coverslips or glass bottom dishes (MatTek Corporation, No. 1.5, 14 mm circular) coated with 5 μg/mL fibronectin (EMD Millipore, MA). Prior to imaging, cells were incubated with Halo JF 646 HaloTag Ligand (#GA1121, Promega, Madison, WI) for 20 min. Imaging was carried out on a Nikon Eclipse Ti-E inverted microscope equipped with a 4-color spinning disk Yokogawa CSU-X1, 405/488/568/647 nm lasers, a Photometrics 95B-illuminated sCMOS camera, and an Apochromat TIRF ×100 NA 1.49 oil lens. Z-stacks with planes separated by 0.4 μm over the whole cell volume were acquired every 1 min. Maximum intensity projections were generated for whole-cell visualization.

### Western blotting

Western blotting was performed using the Protein Electrophoresis and Blotting System (Bio-Rad, Hercules, CA). Briefly, cells were collected, lysed using RIPA buffer (#R0278, Sigma-Aldrich, St. Louis, MO) and resuspended in Laemmli Sample Buffer (Bio-Rad, Hercules, CA). Protein samples (20 µg) were resolved on 8% SDS–PAGE gels and transferred to nitrocellulose membranes (GE Healthcare Life Sciences, Chicago, IL) at 350 mA for 3 h for blotting. Membranes were blocked in 5% nonfat dried milk (Sigma-Aldrich, St. Louis, MO) for 1 h and incubated overnight with the following primary antibodies: anti-KIF5B (1:1000, #ab151558, Abcam), anti-KLC1 (1:500, #19028-1-AP, Proteintech, Rosemont, IL), anti-KLC3 (1:500, #sc-398332, Santa Cruz, Dallas, TX), anti-KIFC3 (1:500, #PA5-30074, Invitrogen, Waltham, MA), anti-α-tubulin (1:1000, #ab18251, Abcam, Cambridge, MA), and anti-GAPDH (1:2000, #sc-32233, Santa Cruz, Dallas, TX). Secondary antibodies conjugated to IRDye 700 and 800 (1:5000, LI-COR Biosciences, Lincoln, NE) were used. Membranes were imaged using an Odyssey Infrared Imaging system (LI-COR Biosciences, Lincoln, NE) or ChemiDoc MP Imaging System (Bio-Rad, Hercules, CA).

### Image Processing

Image analysis was performed using NIS-Elements, ImageJ, and MATLAB. For all fluorescence images presented, whole-image histogram stretching adjustments were made in each channel to optimize brightness and contrast and enhance visibility.

### Quantitative Analyses

Centrosome coordinates were obtained using automatic detection of the intensity maximum of the centrioles in the Imaris program (Bitplane) or ImageJ. For Golgi and ERES segmentation, background levels were determined using intensity histograms of maximum intensity projections. A manual threshold above the background was thereafter implemented to full 3D image stacks in ImageJ. Importantly, the background was determined individually for each live imaging sequence to accommodate the variability of the Golgi and ERES marker expression levels. For fixed images, the background levels were identified per experimental repeat to account for variability of immunostaining intensity. To calculate Golgi-centrosome (GC) distances, the distance from each voxel within the segmented Golgi volume to both centrosomes was measured^3,43^. The mean distance to the nearest centrosome was then calculated to assess Golgi’s association with the proximal centrosome. Similarly, for calculating Golgi-ERES (GE) distances, the distance from each voxel within the segmented volume of ERES was determined, and the mean distance to the Golgi centroid was calculated. All quantitative analyses were computed using MATLAB (MathWorks, Natick, MA).

### Statistics

Unless otherwise specified, statistical significance was determined using Student’s *t*-test, with *P* < 0.05 considered significant. All statistical analyses and graphical representations were performed using GraphPad Prism (GraphPad Software, San Diego, CA).

## Resource availability

### Lead contact

Requests for further information, resources, and reagents should be directed to and will be fulfilled by the lead contact, Dr. Irina Kaverina.

### Materials availability

Reagents and plasmids used in this study will be available from the lead contact upon request.

### Data and code availability

This paper does not report original code. Any additional information required to reanalyze the data reported in this paper is available from the lead contact upon request.

## Acknowledgements

This work was supported by National Institutes of Health (NIH) grants NIGMS MIRA R35GM127098 (to I.K) and R01 DK106228 (to I.K). We thank Dr. Vladimir Gelfand for kindly providing the KIF5B knockout RPE1 cell line for this study. We thank Claire Scott and Tung Hoang for their help in method optimization. We thank all the members of the Kaverina laboratory and the Vanderbilt Mechanobiology & More club for their advice and support.

## Author contributions

Project design and conceptualization: A.V.S. and I.K; experimental work: A.V.S.; data analysis: A.V.S.; figure preparation: A.V.S. and I.K.; manuscript draft: A.V.S and I.K.; project supervision: I.K.; funding acquisition: I.K. Both authors approved the final version of the manuscript and agree on the content and conclusions.

## Supplemental information

Document S1, Supplementary Figures 1-2

Supplementary video 1. A time-lapse imaging sequence of an RPE1 cell synchronized in the G2 phase, stably expressing Golgi (RFP-ST, magenta), centrosome (centrin-GFP, green), and ectopically expressed ERES (halo-sec23, yellow) markers treated with RO3306 (CDK1 inhibitor) for 2 h. Maximum intensity projections of spinning disk confocal stacks are shown. Single-channel inverted grayscale are combined to highlight Golgi and ERES dynamics. Arrows indicate centrosomes. Time, hours: minutes. Scale bar: 10 µm. Corresponds to Figure 1E.

Supplementary video 2. A time-lapse imaging sequence of an RPE1 cell synchronized in the S phase, stably expressing Golgi (RFP-ST, magenta), centrosome (centrin-GFP, green), and ectopically expressed ERES (halo-sec23, yellow) markers treated with RO3306 (CDK1 inhibitor) for 60 min. Maximum intensity projections of spinning disk confocal stacks are shown. Single-channel inverted grayscale are combined to highlight Golgi and ERES dynamics. Arrows indicate centrosomes. Time, hours: minutes. Scale bar: 10 µm. Corresponds to Figure 1H.

Supplementary video 3. A time-lapse imaging sequence of an RPE1 cell synchronized in the G2 phase, stably expressing Golgi (RFP-ST, magenta), centrosome (centrin-GFP, green), and ectopically expressed ERES (halo-sec23, yellow) markers treated with kinesore for 2 h. Maximum intensity projections of spinning disk confocal stacks are shown. Single-channel inverted grayscale are combined to highlight Golgi and ERES dynamics. Arrows indicate centrosomes. Time, hours: minutes. Scale bar: 10 µm. Corresponds to Figure 2A.

Supplementary video 4. A time-lapse imaging sequence of an RPE1 cell synchronized in the G2 phase, stably expressing Golgi (RFP-ST, magenta) and centrosome (centrin-GFP, green) markers treated with kinesore for 2 h, washed, and incubated in kinesore-free medium for 3 h. Maximum intensity projections of spinning disk confocal stacks are shown. A single-channel inverted grayscale is combined to highlight Golgi dynamics. Arrows indicate centrosomes. Time, hours: minutes. Scale bar: 10 µm. Corresponds to Figure 2E.

Supplementary video 5. A time-lapse imaging sequence of an RPE1 cell synchronized in the G2 phase, stably expressing centrosome (centrin-GFP, green) and ectopically expressed ERES (halo-sec23, yellow) markers treated with kinesore for 2 h, washed, and incubated in kinesore-free medium for 3 h. Maximum intensity projections of spinning disk confocal stacks are shown. A single-channel inverted grayscale is combined to highlight ERES dynamics. Arrows indicate centrosomes. Time, hours: minutes. Scale bar: 10 µm. Corresponds to Figure 2I.

Supplementary video 6. A time-lapse imaging sequence of an KIF5B KO RPE1 cell synchronized in the G2 phase, and ectopically expressing Golgi (RFP-ST, magenta), centrosome (centrin-GFP, green), and ERES (halo-sec23, yellow) markers treated with RO3306 for 60 min. Maximum intensity projections of spinning disk confocal stacks are shown. Single-channel inverted grayscale are combined to highlight Golgi and ERES dynamics. Arrows indicate centrosomes. Time, hours: minutes. Scale bar: 10 µm. Corresponds to Figure 3E.

Supplementary video 7. A time-lapse imaging sequence of an KIF5B KO RPE1 cell synchronized in the G2 phase, and ectopically expressing Golgi (RFP-ST, magenta), centrosome (centrin-GFP, green), and ERES (halo-sec23, yellow) markers treated with kinesore for 2 h. Maximum intensity projections of spinning disk confocal stacks are shown. Single-channel inverted grayscale are combined to highlight Golgi and ERES dynamics. Arrows indicate centrosomes. Time, hours: minutes. Scale bar: 10 µm. Corresponds to Figure 3H.

Supplementary video 8. A time-lapse imaging sequence of an RPE1 cell synchronized in the G2 phase, stably expressing Golgi (RFP-ST, magenta), centrosome (centrin-GFP, green), and ectopically expressed ERES (halo-sec23, yellow) markers treated with Ciliobrevin D and RO3306 for 60 min. Maximum intensity projections of spinning disk confocal stacks are shown. Single-channel inverted grayscale are combined to highlight Golgi and ERES dynamics. Arrows indicate centrosomes. Time, hours: minutes. Scale bar: 10 µm. Corresponds to Figure 5A.

Supplementary video 9. A time-lapse imaging sequence of an RPE1 cell synchronized in the G2 phase, stably expressing Golgi (RFP-ST, magenta), centrosome (centrin-GFP, green), and ectopically expressed ERES (halo-sec23, yellow) markers treated with Ciliobrevin D and kinesore for 2 h. Maximum intensity projections of spinning disk confocal stacks are shown. Single-channel inverted grayscale are combined to highlight Golgi and ERES dynamics. Arrows indicate centrosomes. Time, hours: minutes. Scale bar: 10 µm. Corresponds to Figure 5D.

Supplementary video 10. A time-lapse imaging sequence of KIFC3 knockdown RPE1 cell synchronized in the G2 phase, stably expressing Golgi (RFP-ST, magenta), centrosome (centrin-GFP, green), and ectopically expressed ERES (halo-sec23, yellow) markers treated with RO3306 for 60 min. Maximum intensity projections of spinning disk confocal stacks are shown. Single-channel inverted grayscale are combined to highlight Golgi and ERES dynamics. Arrows indicate centrosomes. Time, hours: minutes. Scale bar: 10 µm. Corresponds to Figure 5G.

Supplementary video 11. A time-lapse imaging sequence of KIFC3 knockdown RPE1 cell synchronized in the G2 phase, stably expressing Golgi (RFP-ST, magenta) and ectopically expressed ERES (halo-sec23, yellow) markers treated with kinesore for 2 h. Maximum intensity projections of spinning disk confocal stacks are shown. Single-channel inverted grayscale are combined to highlight Golgi and ERES dynamics. Arrows indicate centrosomes. Time, hours: minutes. Scale bar: 10 µm. Corresponds to Figure 5J.

