## Supplementary File S1 for "Cell cycle-regulated tug-of-war between microtubule motors positions major trafficking organelles"

### Supplementary Figures

A

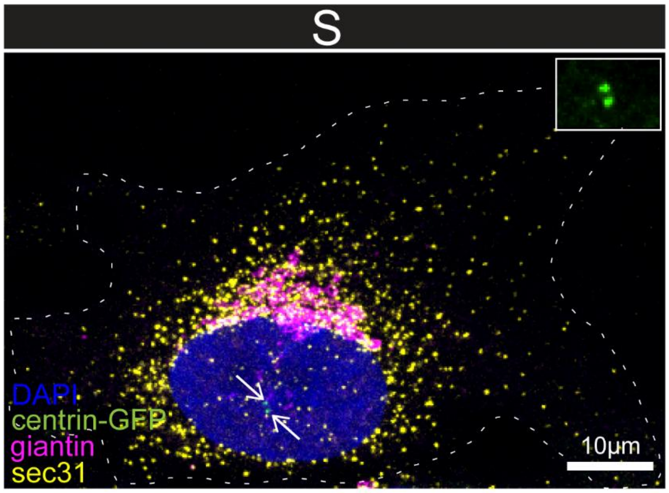


B

| Parameter | Quantification method | |
|---|---|---|
| Golgi-centrosome distance (GC) | 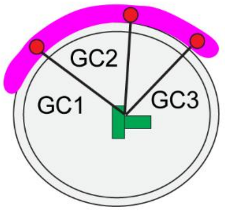 | Average (GC1, GC2, GC3,GC4.....) |
| Golgi-ER exit sites distance (GE) | 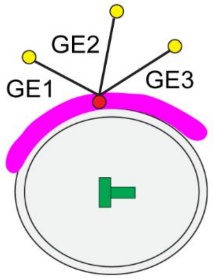 | Average (GE1, GE2, GE3,GE4.....) |

**Supplementary Figure 1.** A. Representative image showing Golgi and ERES positioning in S phase. RPE1 cells stably expressing centrin-GFP (green) were immunostained for DAPI (blue), giantin (magenta), and sec31 (yellow). Image is a maximum intensity projection of NSPARC-acquired Z-stacks. Arrows indicate centrosomes, which are enlarged in solid boxed insets. Scale bar: 10 µm. B. Table summarizing the computational method used to quantify Golgi and ERES positioning (adapted from Frye et al., 2020 and Frye et al., 2023). Golgi-centrosome distance (GC) was calculated by measuring the distance from each voxel of the segmented volume of Golgi to both centrosomes and computing the mean value to represent overall Golgi-centrosome association. Similarly, Golgi-ERES distance (GE) was calculated by measuring the distance from each voxel of segmented volume of ERES to the centroid of the Golgi, with the mean value representing Golgi-ERES spatial association.

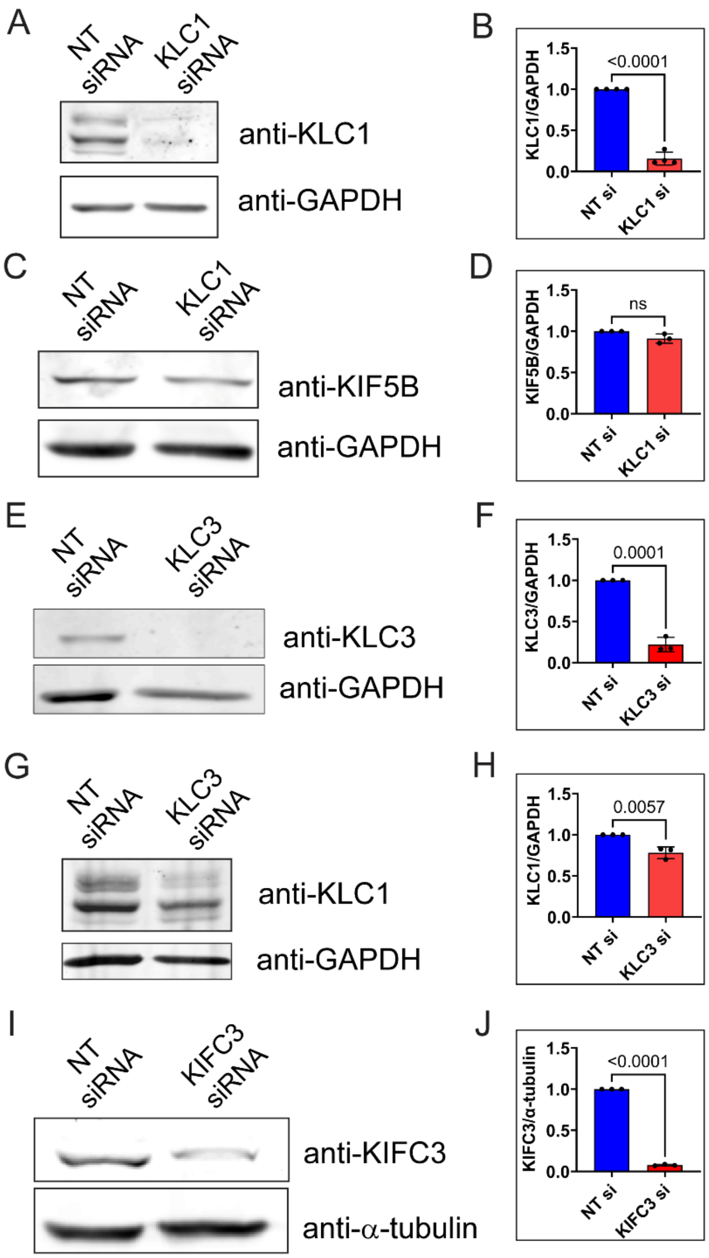

**Supplementary Figure 2.** A. Representative western blot showing KLC1 protein levels following KLC1 siRNA treatment. GAPDH was used as a loading control. B. Quantification of KLC1 protein levels from western blots. Data represent mean ± s.d., n=4. C. Representative western blot showing KIF5B protein levels following KLC1 siRNA treatment. GAPDH was used as a loading control. D. Quantification of KIF5B protein levels from western blots. Data represent mean ± s.d. n=3. E. Representative western blot showing levels of KLC3 protein following KLC3 siRNA treatment. GAPDH was used as a loading control. F. Quantification of KLC3 protein levels from western blots. Data represent mean ± s.d. n=3. G. Representative western blot showing levels of KLC1 protein following KLC3 siRNA treatment. GAPDH was used as a loading control. H. Quantification of KLC1 protein levels from western blots. Data represent mean ± s.d. n=3. I. Representative western blot showing levels of KIFC3 protein following siRNA treatment. α-tubulin was used as a loading control. J. Quantification of KIFC3 protein levels from western blots. Data represent mean ± s.d. n=3.
